# The effect of microsaccades in the primary visual cortex: a two-phase modulation in the absence of visual stimulation

**DOI:** 10.1101/2024.08.12.607606

**Authors:** Yarden Nativ, Tomer Bouhnik, Hamutal Slovin

## Abstract

Our eyes are never still. Even when we attempt to fixate, the visual gaze is never motionless, as we continuously perform miniature oculomotor movements termed as fixational eye movements. The fastest eye movements during the fixation epochs are termed microsaccades (MSs), that are leading to continual motion of the visual input, affecting mainly neurons in the fovea. Yet our vision appears to be stable. To explain this gap, previous studies suggested the existence of an extra-retinal input (ERI) into the visual cortex that can account for the motion and produce visual stability. Here, we investigated the existence of an ERI to V1 fovea in behaving monkeys while they performed spontaneous MSs, during fixation. We used voltage-sensitive dye imaging (VSDI) to measure and characterize at high spatio-temporal resolution the influence of MSs on neural population activity, in the foveal region of the primary visual cortex (V1). In the absence of a visual stimulus, MSs induced a two-phase response modulation: an early suppression transient followed by an enhancement transient. A correlation analysis revealed an increase in neural synchronization around ∼100 ms after MS onset. Next, we investigated the MS effects in the presence of a small visual stimulus, and found that this modulation was different from the non-stimulated condition yet both modulations co-existed in the fovea. Finally, the VSD response to an external motion of the fixation point could not explain the MS modulation. These results support an ERI that may be involved in visual stabilization already at the level of V1.

## Introduction

Our eyes are never still. We scan the surroundings with large and rapid ballistic eye movements termed as saccades (size > 1 deg) which enable us to gather visual information from points of interest in the visual scene. Even when we attempt to fixate, the visual gaze is never entirely motionless, because we continuously perform miniature oculomotor movements termed as fixational eye movements. These tiny eye movements are classified into two main categories: Microsaccades (MSs) and Drift. Microsaccades are fast and small (size < 1 deg) fixational saccades and are considered to be the smaller version of saccades, while Drift is the slow eye motion during the inter-saccadic interval (Martinez-Conde et al., 2004, 2009; Rolfs, 2009; Rucci and Victor, 2015; Ahissar et al., 2016; Krauzlis et al., 2017).

In primates, MSs and saccades enhance visual processing by allowing to flexibly allocate the fovea, a specialized high-acuity zone at the center of the primate’s retina, towards points of interest in the visual scene (Green, 1970; Wurtz, 2008; Poletti et al., 2013; Intoy et al., 2021). However, these fast and ballistic eye movements pose also a substantial challenge for the visual system, because each MS or saccade produces an abrupt and rapid motion of the visual input over the retina. Yet, despite the continual displacement of the visual input on the retina, we don’t perceive these shifts and our visual perception remains stable. Saccadic suppression, which is a decrease in sensitivity to a visual stimulus, was suggested as a mechanism that can mediate stable vision (Volkmann, 1986; Diamond et al., 2000; Burr, 2004; Binda and Morrone, 2018). Recently, an increase in visual thresholds was described also for MSs (MSs suppression; Scholes et al., 2018; Intoy et al., 2021). Moreover, the decrease in sensitivity starts before the MS or saccade onset, suggesting the existence of an extra-retinal input (ERI; for example: an efference copy of the motor command to the oculomotor muscles).

Past neurophysiological studies investigated the effects of saccades and MSs on neuronal activity in the visual cortex of humans and monkeys (for reviews see: Ibbotson and Krekelberg, 2011; Krauzlis et al., 2017; Martinez-Conde et al., 2013; Rolfs, 2009; Ross et al., 2001; Wurtz, 2008). Studies investigating the effects of saccades on neural responses in animlas, reported on neural modulation in various visual cortical areas, comprised of neural suppression, neural enhancement or both (Royal et al., 2006; Bremmer et al., 2009; Idrees et al., 2020; Parker et al., 2023). Several studies investigated the existence of a saccade related ERI into the visual cortex (Royal et al., 2006; Rajkai et al., 2008; Miura and Scanziani, 2022; Niemeyer et al., 2022; Denagamage et al., 2023).

The effects of MSs on neuronal responses in monkeys and humans were also studied in various visual areas, and reported to be early suppression, or late enhancement or a sequence of early suppression followed by an enhancement (Kogan et al., 2008; Meirovithz et al., 2012; Gilad et al., 2014; Troncoso et al., 2015; Gilad et al., 2017; Wu et al., 2022). The effects of MSs in early visual cortex, during visual stimulus presentation showed that at least part of the modulation can be explained by the stimulus displacement on the retina (Meirovithz et al., 2012). Only few studies focused on the involvement of ERI in MSs and investigated the effects of MSs without a visual stimulus in V1 or V2 (Snodderly et al., 2001; Troncoso et al., 2015; Wu et al., 2022).

The possible existence of an ERI into V1, suggests also a modulation in neural synchronization. Yet, only very few studies investigated the effects of neural synchronization around saccades onset (Maldonado et al., 2008; Niemeyer et al., 2022; Denagamage et al., 2023) and MSs onset (Bosman et al., 2009; Meirovithz et al., 2012; Lowet et al., 2018; discussed in: Leopold and Logothetis, 1998; Martinez-Conde et al., 2004). In particular, the relation between the changes in synchronization and MSs in the absence of visual stimuli in early visual cortex were not studied.

Here we used voltage-sensitive dye imaging (VSDI) in monkeys to investigate the effects of MSs on the neural population responses in the foveal region of V1, when the animals fixate on a blank screen (i.e. no visual stimulus except for a tiny fixation point). Past neurophysiological studies did not focus on V1 fovea and fovela, where neurons are expected to be most influenced by the tiny fixational saccades. Moreover, most of these studies focused mainly on MSs effects in the presence of a visual stimulus. Here we applied a different approach: using VSDI, we characterized the influence of MSs on V1 activity at the fovea, in the absence of visual stimulation, except for a tiny fixation point. Our results show that MSs induce a bi-phasic modulation in V1, comprised of suppression followed by enhancement. This bi-phasic modulation was associated with a transient increase of neural synchronization. Additionally, the MS modulation in the presence of a small visual stimulus was different from that in the absence of a visual stimulus. Finally, we compared the neural response evoked by an artificial movement of the fixation point (mimicking the fixation point movement on the retina) to that generated by MSs. Overall, our results reveal robust evidence for ERI related to MSs in V1.

## Methods

### Animals and dataset

Three adult male monkeys (Macaca fascicularis, males, 8-12 Kg; monkeys L, G, B) were used in this study. 9 recording sessions (3 from each monkey) were used for retinotopic mapping of the imaged chamber and defining the V1/V2 border (see additional details in: ’Defining the border between V1 and V2 and retinotopic mapping of V1’). Data analysis on MSs effect was performed on a total of 21 recording sessions from 3 hemispheres of the 3 monkeys (Fig. 1-5). The analysis of MSs modulation in blank trials was done on a total of: 6 recording sessions (∼29 trials per session) from the left hemisphere of monkey G, 6 recording sessions (∼60 trials per session) from the right hemisphere of monkey L and 1 recording session (68 trials) from the right hemisphere of monkey B. In each session we analyzed only correct trials that were carefully inspected for eye movements. A total of 521 MSs were analyzed from the 3 monkeys (Monkey L: 219 MSs, amplitude 0.45±0.04 deg; monkey G: 243 MSs, amplitude: 0.37±0.02 deg (mean±sem); Monkey B: 59 MSs, 0.36±0.02 deg).

**Figure 1:**
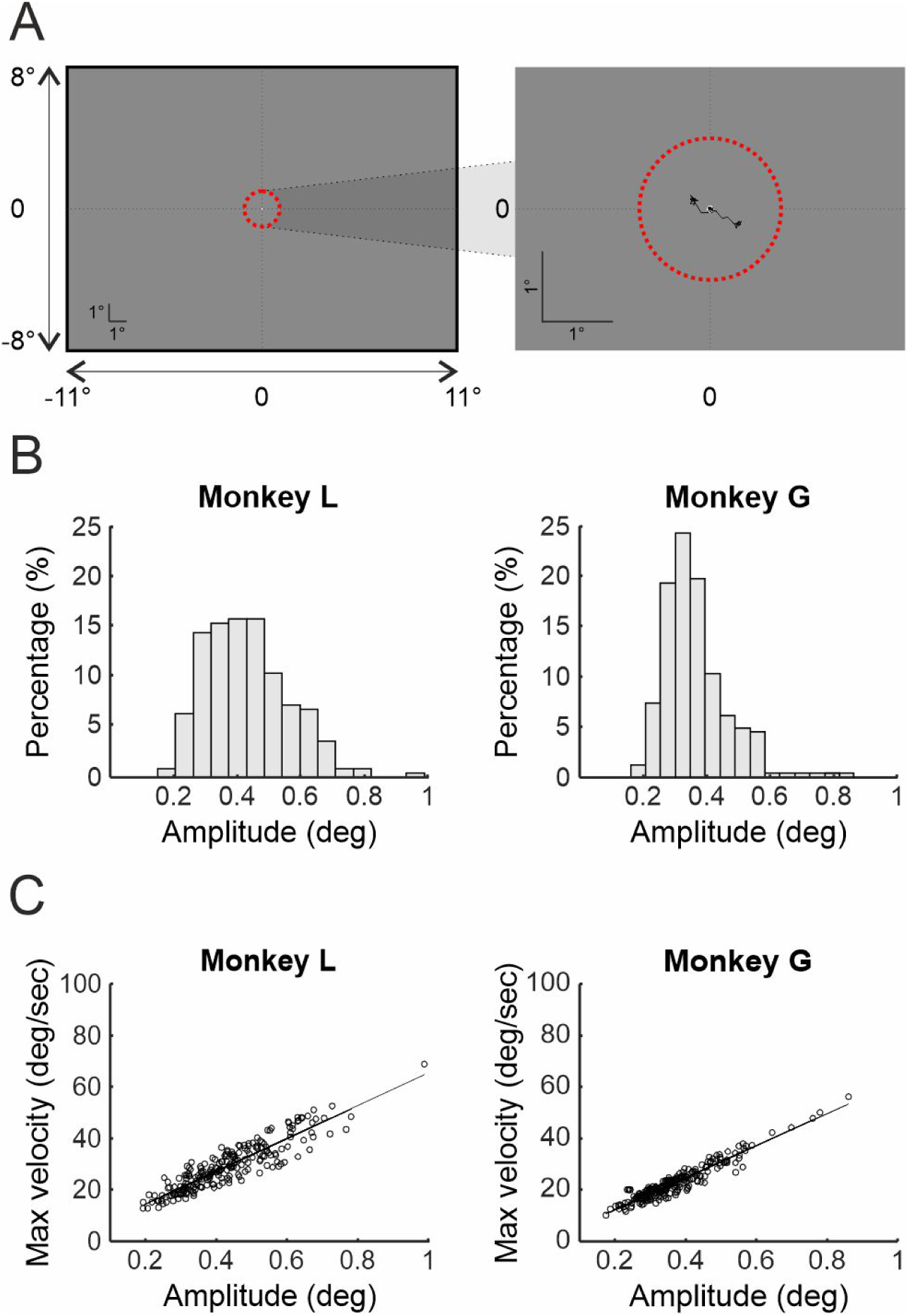
A schematic illustration of the fixation screen and microsaccades (MSs) characteristics. **A**. Left: The monkeys fixated on a small fixation point (0.1-0.2 deg), positioned at the middle of a gray screen (coordinates: 0,0; fixation point shown in scale). The red contour represents the fixation window (radius = 1 deg; not visible to the animals) where the monkeys have performed MSs while maintaining fixation. Dashed vertical and horizontal lines indicate the center position (zero) on each axis. Right: An enlargement of the screen display around the fixation point, with an illustration of two example MSs, one from each monkey. **B.** Distribution histograms of MSs amplitude for each monkey (n=219 monkey L; n=243 monkey G). **C.** The main sequence computed for MSs from each animal: max velocity vs. amplitude. The continuous is line a linear regression of the data. The correlation between maximal velocity and amplitude: r=0.91, p<0.001, monkey L; r=0.96, p<0.001, monkey G.

The analysis on trials with a small visual stimulus (Fig. 4) was done on 50 MSs from 5 sessions (monkey G, amplitude 0.53±0.02 deg). Another set of control experiments of simulated MSs were done in Monkey B and included: 56 trials of artificial FP movement from 3 sessions (30 trials with inward movement of the FP; 26 trials with FP outward movement; Fig. 5). Additional details on the behavioral paradigms are below (see ’Behavioral paradigms and data acquisition’).

### Surgical procedure and voltage-sensitive dye imaging

The surgical procedure and voltage-sensitive dye (VSD) staining have been reported with details, previously (Shoham et al., 1999; Shtoyerman et al., 2000; Arieli et al., 2002; Slovin et al., 2002).

All experimental procedures were carried out according to the NIH guidelines, approved by the Animal Care and Use Guidelines Committee of Bar-Ilan University, and supervised by the Israeli authorities for animal experiments. Briefly, the monkeys were anesthetized, ventilated, and an intravenous catheter was inserted. A head holder and two cranial windows (25 mm, i.d.) were bilaterally placed over the primary visual cortices and cemented to the cranium with dental acrylic cement. After craniotomy, the dura mater was removed, exposing the visual cortex. A thin, transparent artificial dura of silicone was implanted over the visual cortex. Appropriate analgesics and antibiotics were given during surgery and postoperatively. The center of the imaged V1 area for the three animals laid 0.75–1.5 deg below the horizontal meridian and 1–2 deg from the vertical meridian. We stained the cortex with RH-1691 or RH-1838 VSD supplied by Optical Imaging, Israel.

VSDI was performed using the Micam Ultima system based on a sensitive fast camera with up to 10 KHz sampling rate. We used a sampling rate of 10 ms per frame with a spatial resolution of 10,000 pixels where each pixel measures the activity from an area of 170² µm². Each pixel sums the VSD signal from the population activity of a few hundreds of neurons, mainly from the upper layers (2-3) of the cortex. The fluorescence dye signal from each pixel reflects the sum of the membrane potential from all the neuronal elements of that pixel, i.e. a population signal (rather than a single neuron). Thus, the VSD signal reflects both subthreshold and suprathreshold membrane potentials from all neurons. The exposed cortex was illuminated by an epi-illumination stage with appropriate excitation filter (peak transmission 630 nm, width at half-height 10 nm) and a dichroic mirror (DRLP 650), both from Omega Optical. To collect the fluorescence and reject stray excitation light, a barrier postfilter was placed above the dichroic mirror (RG 665, Schott).

### Behavioral paradigms and data acquisition

Two adult male monkeys (monkeys G, L) were trained on a fixation paradigm. Each trial started when a small fixation point (FP, 0.1 deg) appeared in the center of a display screen (coordinates: 0,0) on a gray background (Fig. 1A; note that the FP appears 8-11 deg away from the screen edges). The animals were required to maintain fixation throughout the entire trial duration, that lasted for 4-5s, during which VSD data acquisition (DAQ) was performed. In blank trials, no visual stimulus appeared (except for the FP). In stimulated trials, after a random fixation interval of 3-4s, a small visual stimulus was turned on for a variable duration (small black or white dot; radius: 0.05-0.15 deg; position: 0.75 deg below the horizontal meridian; 0.6 deg from the vertical meridian). During all trials the animals were required to maintain tight fixation within a small fixation window. The animals were rewarded only if the trial was successfully completed. Only correct trials were used for analysis.

Another set of control experiments were performed on a third monkey (monkey B). The animal performed a fixation task (as described for monkey G and L) on a small FP (0.2 deg) that appeared at the center of a gray screen (coordinates: 0,0 deg). In part of the experiments the FP moved to one out of two possible locations: (i) 0.25 deg below and 0.25 deg left to the initial position of the FP (0,0) (ii) 0.2 deg above and 0.2 right to the initial position of the FP (0,0) (Figure 5A). Following a period of 200-300 ms the FP returned to its initial position. In another set of experiments, a small green or white point (0.1 deg) shifted between the above two locations, for a period of 200-300 ms. The spatial shift range of the FP and the small point was 0.3-0.6 deg, thus mimicking typical amplitude size of MSs. Therefore, both control experiments allowed us to simulate an artificial movement of FP, similar to the FP shift on the retina occurring when a MS is generated. Trials were divided into two groups based on the motion direction of the FP (or the point stimulus): (i) the FP (or the point stimulus) moved into the imaged cortical region (inward movement). This simulated a MS away from the imaging chamber, which leads to a spatial shift of the FP towards the chamber (ii) the FP (or the point stimulus) moved outside the imaging chamber. This simulated an inward MS, that is directed towards the chamber, which leads to a spatial shift of the FP away from the chamber. In both experiments the animal was required to maintain fixation and was rewarded only if the trial was successfully completed.

Data acquisition was done using one of the following setups: (i) two linked computers controlled the behavioral task, visual stimulation, data acquisition, and the recording of the monkey’s behavior (CORTEX software package). We used a combination of imaging system (Micam Ultima) and the NIMH-Cortex software package. This system was equipped with a PCI-DAS 1602/12 card to control the behavioral task and data acquisition. (ii) a computer installed with MonkeyLogic (ML) software and equipped with a NI card to control the behavioral task and the data acquisition. We used a combination of ML with the imaging system (Micam Ultima). For both setups, behavioral and neuronal data from single trials was saved in separate files to enable single trial analysis. Additional details on the protocol for VSDI DAQ has been described in detail elsewhere (Slovin et al., 2002).

### Eye position recording and detection of MSs

Eye position was monitored by an infrared eye tracker (Dr. Bouis; Bach et al., 1983), sampled at 1 kHz, recorded at 250-1000 Hz. During the behavioral task, the animals were required to maintain fixation on a small fixation point and typically made a few MSs per second (Fig. 1A). To detect the MSs on each trial, we implemented an algorithm for MSs detection on the monkeys’ eye position data (Engbert and Mergenthaler, 2006). MSs were detected in 2D velocity space of the eye position, using thresholds for peak velocity (in units of median-based standard deviations SDs) and a minimal MS duration. This produced an elliptical threshold in the 2D velocity space (3.5-6 MAD from median velocity). Additionally, the threshold for minimal MS duration was set to 7-12 ms and a minimal 50 ms interval was set between successive MSs. MSs were also defined according to their kinematic properties (amplitude < 1 deg and maximal velocity < 100 deg/s; Engbert and Mergenthaler, 2006; Meirovithz et al., 2012). To minimize noise contamination in the dataset, we set a lower limit for MS amplitude (0.1 deg). The reliability of the algorithm for MSs detection in our data was already confirmed in previous VSD studies (Meirovithz et al., 2012; Gilad et al., 2014).

To compute the MS start point (MS onset) and end point (MS landing), we used a modified algorithm based on previous studies (Hafed et al., 2009; Chen and Hafed, 2013). The eye position traces were filtered using a low-pass filter (3rd-order Savitzky-Golay filter, 25 ms window) and the velocity was re-computed (Cherici et al., 2012). Then, a narrow time window (± 30 ms) was defined around each detected MS, and all velocity points above a threshold (exceeding 3-8 deg/sec; set for each animal based on the data SNR) were used for acceleration analysis. Next, a threshold of acceleration (1-2 MAD from the median acceleration) was used to determine the MS onset and MS landing. MS direction was computed from MS onset to MS landing and the MS amplitude was defined as max amplitude.

In addition, we plotted the amplitude histograms as well as computed the main sequence for all MSs (i.e., the relation between peak velocity and amplitude; Fig. 1B,C). MSs that were 3 STD away from the mean distribution of the main sequence were removed. In addition, we visualized the eye position traces and verified each detected MS.

### Defining the border between V1 and V2 and retinotopic mapping of V1

A total of 9 recording sessions were used for retinotopic mapping and defining the V1/V2 border. In each monkey, we identified V1 and V2 regions in the imaged cortical area using optical imaging of intrinsic signals (Shtoyerman et al., 2000). Ocular dominance and orientation maps were used to identify the border between V1 and V2 areas in each animal. To obtain the retinotopic maps in V1 we used optical imaging of intrinsic signals combined with visual stimulation of horizontal or vertical bars and VSDI combined with visual stimulation of small points (Shtoyerman et al., 2000; Slovin et al., 2002). Based on the empirical retinotopic maps and using a 2D analytical retinotopic model (Ayzenshtat et al. 2012), we could approximately define the visual field eccentricities in the imaged chamber: 0.25-3 deg and 0.2-2.5 deg for monkeys L and G respectively. According to previous studies, the fovea region is defined from 0 to 2.5 deg eccentricity, and parafovea from 2.5 to 5 deg eccentricity (Green, 1970; Moiseenko et al., 2018; Chen et al., 2019). We then defined the foveal and parafoveal regions in the monkeys imaged V1, for each animal (see Fig. 2A,E maps=-60 ms). Finally, we also marked the 0.75 deg eccentricity on the imaged V1, and termed it as the central fovea.

**Figure 2:**
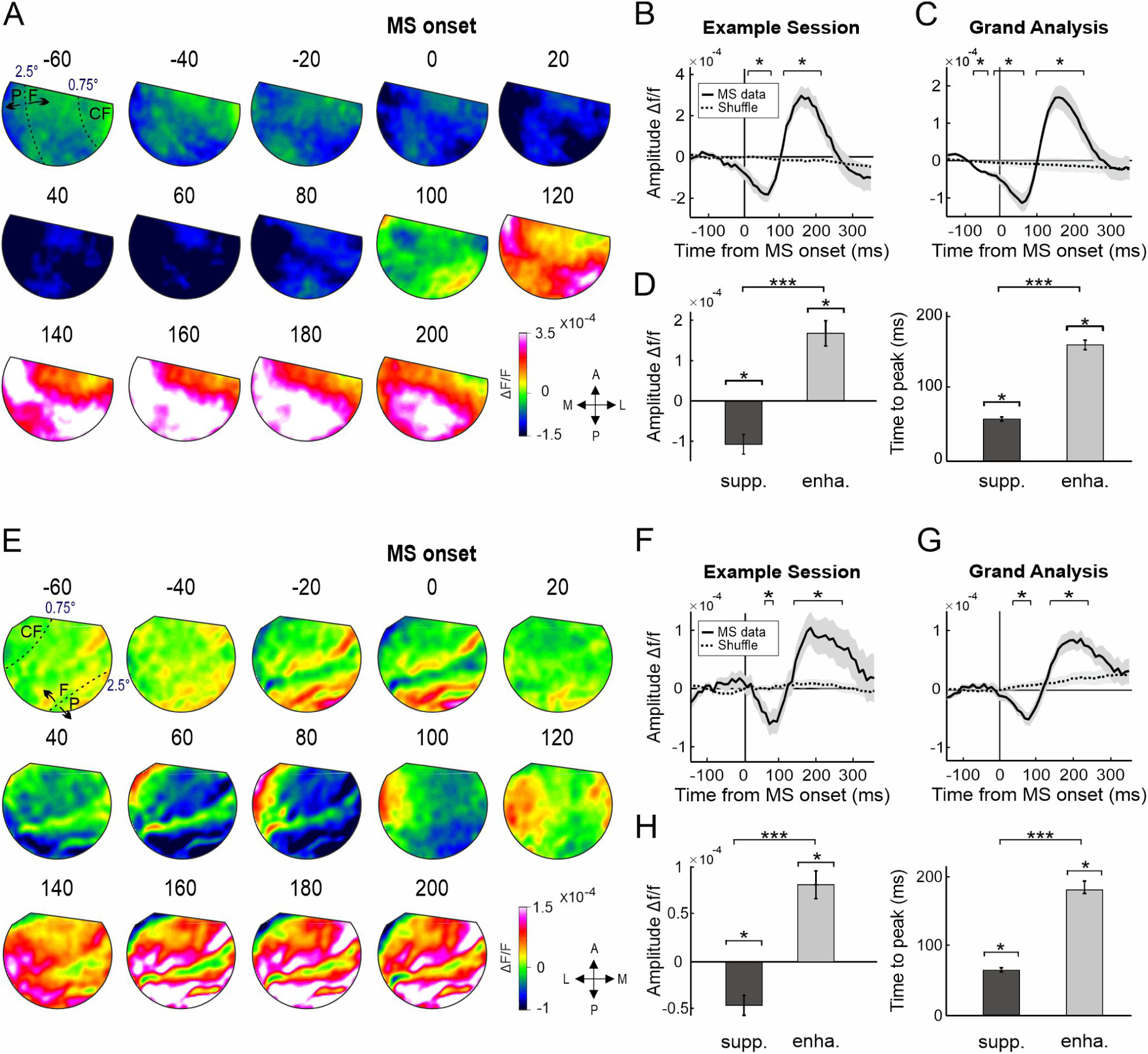
The effects of microsaccades on population responses in blank trials. **(A)** A sequence of VSD maps in V1 aligned on MS onset (t=0; mean across MSs, n=34; monkey L), an example session. Each VSD map is averaged over 20 ms and color coded (blue and red denote suppressed and enhanced activity relative to the baseline). The baseline response (150 ms to 50 ms before MS onset) was subtracted for each pixel as well as the shuffled response (see Materials and Methods). Maps were low-pass filtered for visualization purposes only. The map at t=-60 ms shows the approximate eccentricity lines of 0.75 deg and 2.5 deg, denoting the regions of the fovea (F), parafovea (P) and central fovea (CF) in the imaged V1 (see Materials and Methods). **(B)** Time course of the VSD signal in V1 (mean over all pixels) for the example session in A. The effect of MSs appears in a continuous curve and the shuffle data in a dashed curve (see Materials and Methods). The shaded area represents ±1SEM across trials or shuffle data. **(C)** Grand analysis of MSs modulation in monkey L (n=6 sessions): time course of the VSD signal in V1. Shaded area represents ±1SEM across sessions. The top bars in B and C mark the time points with significant difference between MS modulation and shuffled data (p value range: < 0.001 to < 0.05). **(D)** Left: Suppression and enhancement peak amplitude (mean across all sessions), grand analysis. Right: Time to peak for the MS suppression (supp.) and enhancement (enha.), mean across all sessions, grand analysis. Error bars represent ±1SEM across sessions. * p > 0.05; *** p > 0.001 Wilcoxon rank sum. E-H: as in A-D, but for monkey G. E-F: n=60 MSs; G-H: grand analysis, n=6 sessions.

### Basic VSDI analysis and aligning the VSD signal on MS onset

Basic VSD data processing was described in details previously (Slovin et al., 2002; Ayzenshtat et al., 2010). Briefly, this consisted of choosing pixels with minimal threshold fluorescence, then normalizing each pixel to its baseline fluorescence level and, finally, subtracting the average blank condition to remove the heartbeat artifact. This processing removes, in an unbiased manner, most of the slow fluctuations originating from heartbeat artifact or dye bleaching within a trial (Shoham et al., 1999). These steps were schematically illustrated and explained in Fig. S12 at Ayzenshtat et al., 2010. In addition, we removed pixels in the vicinity of large blood vessels and pixels outside the margins of the imaged cortical area. Single trials that deviated more than 2.5 STDs from the mean (across trials) response, were furthered removed from analysis to avoid noise contamination (e.g. animal motion artifact). VSD maps were low-pass filtered with a 2D Gaussian filter (sigma = 2 pixels) for visualization purposes only.

To analyze the effect of MSs on the VSD signal, we analyzed MSs occurring at 200-1300 ms from onset of VSD DAQ in the blank condition and for the stimulated condition MSs were analyzed in a time window of 100 ms after stimulus onset until 50 ms after stimulus offset. Next, the VSD signal was aligned on MS onset, using a 500 ms window: 150 ms before MS and 340 ms after MS onset. Next, we subtracted from each pixel the baseline activity (mean response at 150 to 50 ms before MS onset) and compared the VSD response to the MS shuffled condition (see ’Statistical analysis and computing the MS shuffled data’). The peak response in MS modulation (negative or positive peak) was computed by averaging the VSD signal within a 30 ms window around the peak response time. Matlab software was used for all calculations and statistical analyses.

### Correlation analysis

To investigate the effect of MSs on synchronization between neural population, we computed the correlation between V1 pixels, for each MS (Meirovithz et al., 2010, 2012). Using a sliding window of 100 ms, a Pearson correlation coefficient (r) was computed between the VSD response in each V1 pixel (Pi) and all the other pixels in the imaged V1 area, during a MS. This computation resulted in a vector of correlation for pixel (Pi). Next, we computed the mean correlation over this vector and the value was assigned to pixel (Pi). This procedure was repeated for each pixel in V1, which resulted in a correlation map for each single time window (Equation 1). The computation of correlation maps over sequential time windows resulted in a sequence of correlation maps for each MS. Finally, the correlation maps were averaged across MSs within a single session (Fig. 3A). In summary, the correlation value within each pixel in the map, reflects the synchronization between pixel (Pi) and all other pixels in V1. Positive, zero and negative values indicate increased synchronization, no synchronization and de-synchronization between V1 pixels, accordingly. To evaluate the statistical significance, the correlation maps were computed for shuffled data (see ’Statistical analysis and computing the MS shuffled data’) and compared with the original MS data.

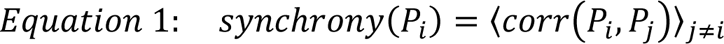

Where *corr* is the Pearson correlation coefficient (r) between pixel i (Pi) and any other pixel in V1, defined as pixel j (Pj). The correlation is then averaged across all possible pairs of P*i* and Pj for a time window of 100 ms. The effect of MSs on synchronization was quantified by computing the mean correlation value across V1 pixels, in the correlation maps. This yielded a time course of correlation (Fig. 3B,C) that was compared with the MS shuffled data. The time points of correlation analysis denote the midpoint of the sliding window (e.g., t = 0 ms shows the correlation value at time window of -50 ms (before MS onset) to 50 ms (after MS onset)). Similar correlation results were obtained using shorter or longer time windows.

**Figure 3:**
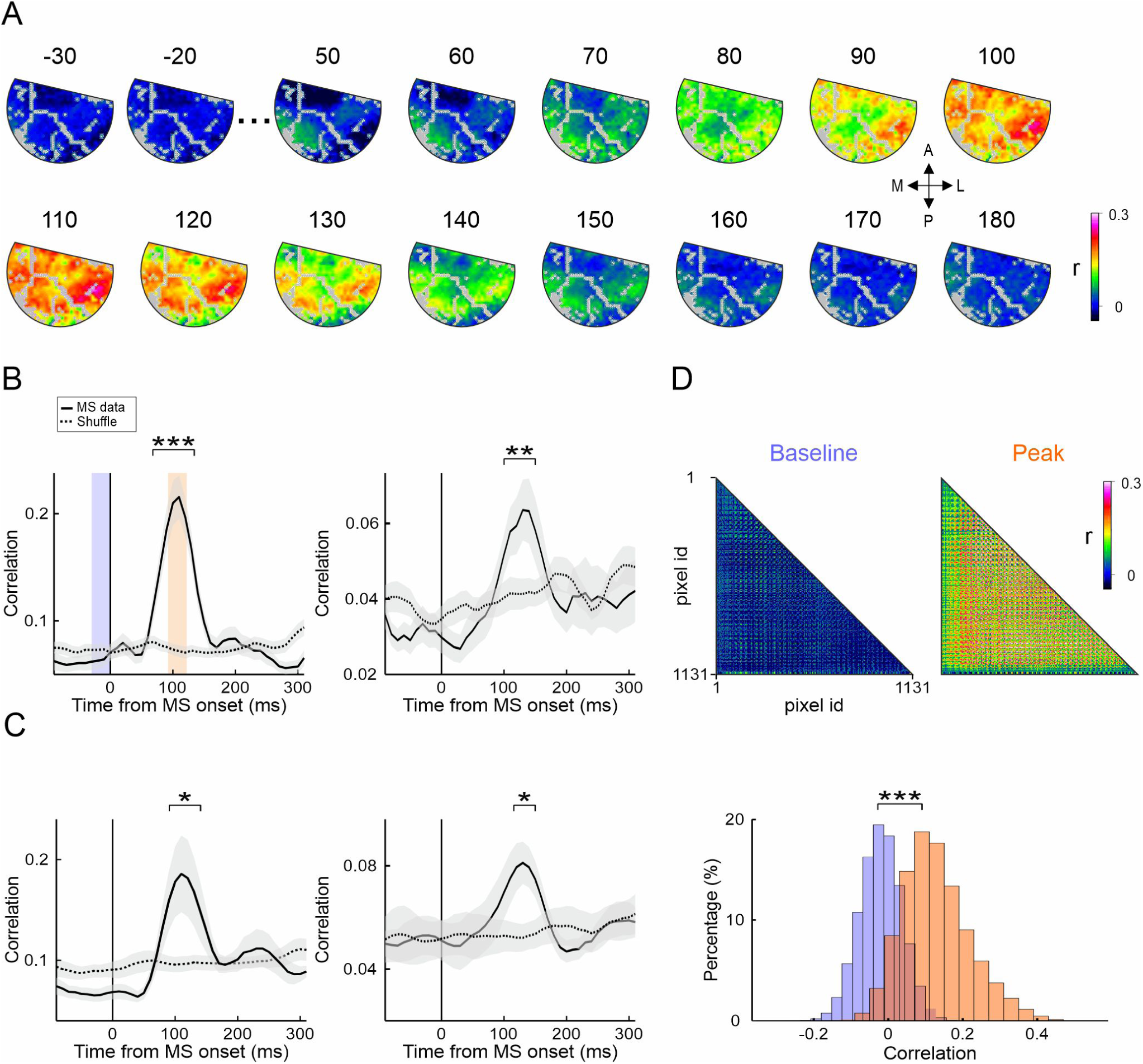
Microsaccades induce a transient increase of synchronization. **(A)** A sequence of mean correlation maps for all V1 pixels in monkey L, with a 100 ms sliding window, example session (n=42 MSs). The value of each pixel in each map represents the correlation (r) between the pixel and all the other pixels in V1 at that time window. The correlations were averaged across the MSs of the same session and the shuffle correlation was subtracted. The gray pixels denote blood vessels that were removed from analysis. **(B)** Time courses of correlation averaged over all V1 pixels in example sessions for monkey L (left; same data as in A; n=42 MSs) and monkey G (right; n=66 MSs). The continuous line represents MSs data, and dashed line the shuffle data. Shaded areas represent ±1 SEM and the top bar with asterisk depicts the time window with a significant difference between the correlation for MSs and shuffle data (Wilcoxon rank sum, * p > 0.05; ** p > 0.01; *** p > 0.001). Shaded blue and orange bars denote the time window used for the single pair-wise correlation matrices in (D). **(C)** Grand analysis of correlation time courses for monkey L (left; n=6 sessions) and monkey G (right; n=6 sessions). **(D)** Top: pair-wise correlation matrices (same example session as in A). The matrices were computed for baseline period (correlations were averaged over 30 ms before MS onset) and peak correlation period (correlations were averaged over 100 to 120 ms after MS onset). The shuffle correlation was subtracted. Bottom: Histograms of the pair-wise correlation (r) from each matrix (***, p<0.001).

Finally, Figure 3D shows the correlation matrix between all single pairs of pixels (rather than averaging across of pairs for a single pixel, as shown in Fig 3A and Equation no. 1; shown for two time windows).

### Analysis of MSs effects in trials with a small visual stimulus

To investigate the MS effects in trials with a small visual stimulus (Fig. 4; a black or white circle, radius = 0.05-0.15 deg; position: 0.75 deg below the horizontal meridian and 0.6 deg from the vertical meridian), we selected trials with the following criteria: 1) The VSD maps showed a clear patch of activation evoked by the stimulus. 2) The properties of the MSs (typically amplitude of 0.2-0.7 deg) enables to visualize the neural activation displacement over the imaged V1 area. Next, we fitted an elliptical ROI to the stimulus evoked activity in each trial with MS (as previously done in Meirovithz et al. 2012). We then computed the VSD signal using the ROIs and time courses were smoothed (with a window size of 3 time points). We defined three ROIs: (i) a ROI fitted to the stimulus evoked response before MS onset, pre-MS ROI (Fig. 4C, left top) (ii) a ROI fitted to the stimulus evoked response after MS landing, pot-MS ROI (Fig. 4C, left middle) (iii) a ROI of V1 area excluding the stimulus evoked response, as defined in (i) and (ii). This ROI was termed as the Non-stimulated ROI (Fig. 4C, left bottom). We verified no overlap between the different ROIs.

**Figure 4:**
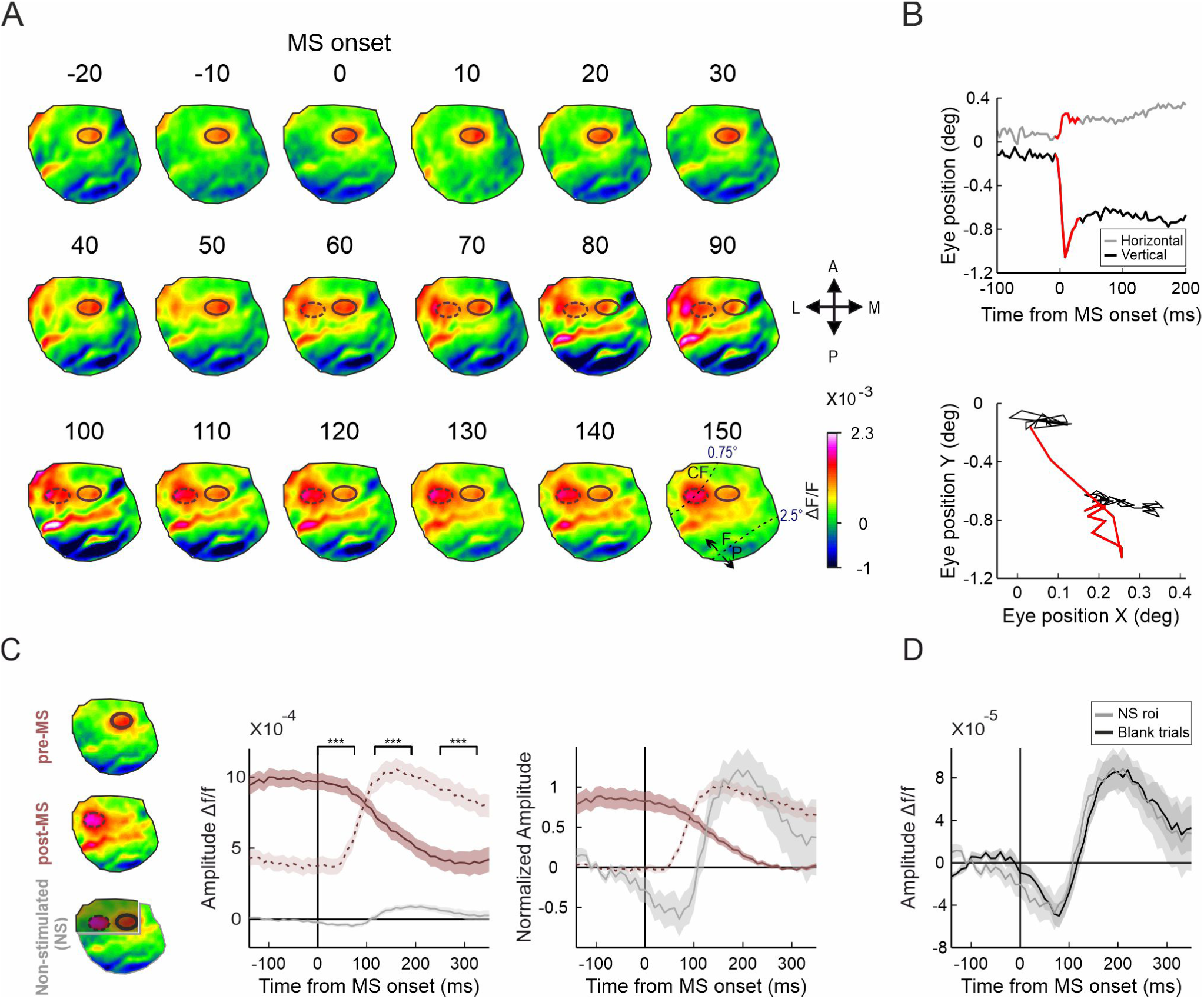
The effects of microsaccades in trials with a small visual stimulus and the comparison to non-stimulated trials. **(A)** A sequence of VSD maps aligned on MS onset in the presence of a small visual stimulus (black dot; position: 0.75 deg below the horizontal meridian; 0.6 deg from the vertical meridian; size: r=0.1 deg), an example trial. The contour lines depict an ellipse fit for two activation patches: prior to the MS (pre-MS; continuous line) and after MS landing (post-MS; dashed line). The map at t=150 ms shows the approximate eccentricity lines of 0.75 deg and 2.5 deg, denoting the regions of the fovea (F), parafovea (P) and central fovea (CF) in the imaged V1 (see Materials and Methods). **(B)** Eye position traces of the example trial in A and the detected MS (in red). Top: the horizontal and vertical eye position in time. Bottom: the eye position in 2D space. **(C)** Left: Maps of the three analyzed ROIs in V1 of the example trial in A: pre-MS ROI, post-MS ROI and the rest of the imaged region in V1, i.e. the non-stimulated (NS) ROI. Middle: mean VSD signal in each ROI (n=50 MSs). Baseline activity was subtracted from the VSD signal in the NS-ROI. Shaded areas represent ±1 SEM across MSs. Top bars with asterisk depict the time points with significant difference between the NS-ROI modulation to the post-MS ROI and pre-MS ROI modulations (Wilcoxon rank sum test; ***, p<0.001). Right: same data as in C middle, but rescaled for comparison of the TC dynamics. **(D)** Time courses of the VSD signal in the NS-ROI averaged across all stimulated trials in comparison to the MS modulation of the grand analysis, in blank trials (Fig. 2G). The correlation coefficient (r) between the two responses is 0.96 (p < 0.001).

### Statistical analysis and computing the MS shuffled data

To evaluate statistical significance, we used nonparametric statistical tests, signed rank test to compare a population’s median to zero and Wilcoxon ranksum test to compare between two medians from two populations. To assess the statistical significance of the MSs modulation in the VSD signal, we created MSs shuffled data. In each recording session, data was pooled across all trials and the original MSs onset times were dissociated from the matched VSD trials and then randomly shuffled over the VSD trials (Meirovithz et al., 2012). This approach preserved the statistical distribution of MSs onset times relative to the VSD signal. Next, we computed the VSD response aligned on the shuffled MS data, for each session and computed the VSD response over ∼1000 shuffled MSs. Similar results were obtained when computing random times of MSs in the VSD trials. To compute the shuffled condition for the correlation analysis, we used a random time for MS onsets relative to the VSD trials (the random times for shuffled data were at least 100 ms shifted from the real data). For each session, we computed ∼100 shuffled MSs.

## Results

Population responses were recorded from V1 area in the right and left hemispheres of two monkeys while they performed a fixation task. The animals were maintaining fixation, i.e. holding their eye gaze on a small fixation point that appeared at the center of a visual display with a gray background (blank trials, see Materials and methods; Figure 1A left). VSDI was performed in the foveal and parafoveal regions of V1 (lower visual field, approximate eccentricity: 0.25-3 deg and 0.2-2.5 deg for monkeys L and G respectively; see Materials and methods). While the animals were maintaining fixation, they performed spontaneous miniature fixational saccades i.e. microsaccades (MSs; Fig. 1A right). The mean MSs amplitude was 0.45±0.04 (mean±SEM); and 0.37±0.02 for monkeys L and G respectively (Fig. 1B) and the relation between MSs amplitude and maximal velocity confirmed the main sequence (Fig. 1C; r=0.91, p<0.001 for monkey L; r=0.96, p<0.001 for monkey G).

To measure the neural activity evoked by MSs, we used VSDI at high spatial (mesoscale, 170^2^ µm^2^/pixel) and temporal resolution (100Hz) from a V1 area, spanning an area of 14-16 mm in diameter. The fluorescence dye signal from each pixel reflects the sum of the membrane potential from all the neuronal elements of that pixel, i.e. a population signal (rather than the response recorded from a single neuron). Thus, the VSD signal reflects the membrane potential from subthreshold activity (i.e. synaptic potentials) to suprathreshold activity (i.e. spiking activity; see Ayzenshtat et al., 2010; Grinvald and Hildesheim, 2004; Jancke et al., 2004). The sensitivity to the subthreshold activity means that even subtle changes in membrane potential can be detected using VSDI. A main advantage of VSDI in this work, is the ability to measure the activation patterns evoked by MSs, in large parts of the foveal representation in V1, the main region to be influenced by these small fixational saccades (size < 1 deg). In addition, we utilized the VSDI to investigate the MSs effects in the absence visual stimuli, and the relation to MS modulation in the presence of a small visual stimulus, which activated only part of the imaged V1 area.

### Microsaccades induce a bi-phasic modulation in V1 activity

Figure 2 shows the VSD maps, i.e. population response maps in V1 of the two monkeys, aligned on MSs onset in blank trials (no visual stimulus except for the small fixation point). Figure 2A,E depicts a time sequence of V1 VSD maps (20 ms apart) aligned on MSs onset, from two example sessions in monkeys L and G. The first map of each example session illustrates the approximate regions of the fovea for each animal. Negative and positive times denote the neural responses before and after the MS onset. The VSD maps show a bi-phasic response that appears over most V1 area: following MS onset (t=0) there is an early suppression transient (dark-blue pixels denote negative response; ∼0-80 ms and 40-80 ms after MS onset, for monkey L and G respectively) that is followed by an enhancement transient (∼140-200 ms after MS onset; pink-white pixels denote positive response). To quantify this, we computed the time course (TC) of the VSD signal by averaging over all V1 pixels in the example sessions. Figure 2B,F show the VSD TCs in each of the example sessions which reveal a bi-phasic modulation comprised of an early suppression (negative peak at t=70 ms after MS onset for both monkeys) followed by a late enhancement (positive peak at t=160 and 180 ms after MS onset, for monkeys L and G respectively). The negative peak response amplitude of suppression is -1.68±0.35×10^-4^ and -0.51±0.2×10^-4^ ΔF/F (mean ± SEM over MSs) for monkey L and G respectively. The peak response amplitude of enhancement is 2.81±0.48×10^-4^ and 1.0±0.27×10^-4^ ΔF/F for monkey L and G respectively. Both the negative and positive peak responses were significantly different from the shuffled data (p value range: <0.001 to < 0.05; Wilcoxon ranksum test; see Material and Methods).

Next, we computed the grand analysis, i.e. across all imaging sessions, for each animal. The MSs modulation was averaged over all sessions (n=6 for each monkey). Figure 2C,G depicts the TCs in V1, confirming a bi-phasic modulation for both animals and Fig. 2D,H shows the quantification of this modulation. Across sessions, the suppression transient reached a negative peak at 60±2.58 and 73.3±3.33 ms (mean±SEM across sessions) after MS onset for monkey L and G respectively. The enhancement transient reached its peak value at 165.7±6.71 ms and 206.7±10.22 for monkey L and G respectively (Fig. 2D,H right). The negative peak response of the suppression had an amplitude of -1.08±0.24×10^-4^ and -0.46±0.11×10^-4^ ΔF/F for monkey L and G respectively, and peak response amplitude of the enhancement was 1.68±0.31×10^-4^ and 0.82±0.15×10^-4^ ΔF/F for monkey L and G respectively (Fig. 2D,H left; p<0.05 Wilcoxon signed-rank, for significant difference from zero; p<0.005, Wilcoxon rank sum for significant difference between negative and positive peaks).

In summary, the results suggest a clear bi-phasic modulation in V1 response following MSs in blank trials (68-73% of V1 pixels across all recording session showed a significant bi-phasic modulation). Notably, the first phase of the modulation is suppression, which is different from the typical increase in V1 population response to visual stimulus onset (or following a spatial shift of a small stimulus; see Slovin et al., 2002; Meirovithz et al., 2012). In addition, the very small fixation point was located most of the time at the center of the foveola, that is outside the imaging chamber. Thus, we anticipated only minimal effects due to possible visual stimulation of the fixation point itself on the imaged population responses within the optical chamber.

### Microssacdes induce a wide spread, transient increase of synchronization in V1

The analyzed MSs were performed over a homogeneous blank screen and in the absence of a visual stimulus. This suggests that the MS neural modulation is not caused by a visual stimulation, but may reflect a common neuronal input into V1 that can induce changes in neuronal correlation and drives the modulation. To investigate this, we computed the zero time-lag correlation (Pearson correlation coefficient, r) between V1 pixels for each MS (i.e. at the single trial level), using a sliding window of 100 ms (see Materials and Methods). The results of this analysis are correlation maps (i.e. synchronization maps; averaged across MSs), where the value in each pixel indicates the mean correlation between this pixel and all other V1 pixels. Thus, the maps sequence shows the temporal evolvement of correlation between all V1 pixels. An example session with a time sequence of correlation maps is shown in Fig. 3A: the dark blue colors denote lower correlation values and warmer colors denote higher correlation values. The maps show that, around t=80-130 ms (time denotes mid correlation windows) after MS onset, there is an increase in correlation, i.e. increase in synchronization, that is wide spread over V1. The time window of increased synchronization overlaps with the suppression phase of the MSs modulation and the initial part of the enhancement phase (similar results were obtained using shorter or longer time windows; see Materials and Methods).

Next, we plotted the TC of synchronization by computing the mean correlation value for all V1 pixels, in the correlation maps. Figure 3B shows the TCs of correlation from two example sessions of the two monkeys. The correlation for monkey L peaked at t=110 ms after MS onset with a correlation value of 0.22±0.02 and for monkey G at t=130 ms after MS onset with a value of 0.06±0.01, both are significantly different from the shuffled correlation data (see Materials and Methods; Wilcoxon ranksum p<0.001 and p<0.01 for monkey L and G respectively). The correlation analysis in Fig. 3C shows the grand average (mean across all sessions) with similar results to the example sessions: peak correlation amplitude of 0.19±0.04 at t=110 ms from MS onset for monkey L and peak correlation amplitude of 0.08±0.01 at t=130 ms for monkey G (peak correlations are significantly different from shuffled correlation data, p<0.05, Wilcoxon ranksum).

Next, we wanted to test whether the single pair-wise correlation matrix (rather than the mean correlation between a pixel and all V1 pixels; see Materials and Methods) also shows increased synchronization after MS onset. Figure 3D shows the pair-wise correlation matrix for the same example session as in Fig. 3A. Figure 3D top left shows the pair-wise correlation matrix at the baseline period (30 ms before MS onset) and Fig. 3D top right shows the pair-wise correlation matrix at the peak TC of correlation (100 to 120 ms after MS onset; see Fig. 3B for highlighted time windows). Figure 3D bottom shows the distribution histograms of the correlation values in the two time windows and there is a significant difference between the baseline correlation and peak correlation (Wilcoxon rank sum, p<0.001). In summary, the correlation analysis (correlation maps or matrix of single pair-wise correlations) revealed a transient increase in synchronization that was wide spread in V1 following MSs. This result further supports the existence of a driving input into V1 that can generates a transient synchronization, aligned on MSs onset.

### MS modulation in visually stimulated trials and the relation to MS modulation in blank trials

The MS modulation we found appears in blank trials, i.e. in the absence of visual stimulation (except for the small fixation point). Therefore, it is unclear whether this modulation exists in the presence of a visual stimulus and how it relates to the MS modulation reported in the presence of a visual stimulus (Meirovithz et al. 2012). To investigate this, we analyzed trials where the animal was fixating and a small visual stimulus that was retinotopically mapped to the imaged V1 area, appeared over the screen (Fig. 4 and see Material and Methods). The small visual stimulus (position: 0.75 deg below the horizontal meridian; 0.6 deg from the vertical meridian; size: 0.05-0.15 deg) activated only a small part of the imaged V1 area, which enabled us to measure the MS modulation in V1 regions that were directly activated by the visual stimulus and V1 regions that were not activated by it, as in the blank trials.

Figure 4A depicts a sequence of VSD maps aligned on MS onset (t=0), from an example trial where a small visual stimulus was turned on (prior to the MS; the eye position traces of the MS are depicted in Fig. 4B top). The maps show a clear patch of activation before MS onset (t=0) that is corresponding to the stimulus evoked response (VSD maps at -20 to 0). We defined two elliptical ROIs: the neural activation region evoked by the stimulus before the MS onset (pre-MS; ROI; continuous line; see Fig. 4C left-top), and the activation region following the MS landing (post-MS ROI; dashed line; see Fig. 4C left-middle). The eye position traces indicated that the MS was directed toward the visual stimulus, and therefore the stimulus shifted towards the center of the fovea (Fig. 4B bottom). Thus, the VSD maps show that following a MS there is a gradual shift of the stimulus evoked response in the fovea, towards more lateral regions in V1, i.e. towards the center of the fovea (Fig. 4A; VSD maps from 50-150 after MS onset). The last map in Fig. 4A illustrates this shift towards the center of fovea, where the post-MS ROI is shifted closer to the eccentricity line of 0.75 deg.

Next, we wanted to investigate whether the MS modulation in the blank trials, exists also in trials with visual stimulation. We therefore defined the V1 region that was not activated by the visual stimulus as the non-stimulated (NS) ROI (Fig. 4C left-bottom). The NS-ROI thus reflects the “blank” state in the stimulated trials. Figure 4C middle shows the mean VSD TC of response over all stimulated trials (n=50 MSs, 6 sessions) in the pre-MS ROI, post-MS ROI and NS-ROI. As we previously reported (Meirovithz et al., 2012) the population response aligned on MS onset in the pre-MS and post-MS ROIs reflects the stimulus shift over the retinotopic map in V1, as dictated by the shift of the stimulus over the retina. The population response in the pre-MS shows a decrease in activity (continuous curve) while at the post-ROI (dashed curve) there is an increase of response following the MS. The VSD response in the pre-MS ROI decreased from 9.74±0.6×10^-4^ ΔF/F (mean activity over -70 to 0 ms prior to MS onset) to 4.55±0.7×10^-4^ ΔF/F (mean activity over 200-270 ms after MS onset; Wilcoxon rank sum p<0.001, for difference between pre and post MS onset). The VSD response in the post-MS ROI increased from 3.73±0.55×10^-4^ ΔF/F (mean activity over -70 to 0 ms prior to MS onset) to 10±0.76×10^-4^ ΔF/F at peak response (mean activity over 120-190 ms after MS onset; Wilcoxon rank sum p<0.001; for difference between response at pre and post MS onset).

Interestingly, the VSD signal in the NS-ROI shows a bi-phasic modulation aligned on MS onset, and notably, this modulation is ∼ 10 times smaller than that induced by the visual stimulation (Fig. 3C middle, gray curve). There is a significant difference between the VSD response in the NS-ROI and the pre-MS and post-MS ROIs responses in time windows that are corresponding to the suppression phase (0-70 ms after MS onset, Wilcoxon ranksum p<0.001), enhancement phase (120-190 ms, Wilcoxon ranksum p<0.001) and later times 250-320 ms (Wilcoxon ranksum p<0.001). To compare the response dynamics of the MS modulation in all three ROIs we normalized the VSD response (to overcome the large response amplitude difference). Figure 4C right shows the comparison of the normalized VSD response in all three ROIs, and there is a clear temporal difference between the dynamics of the TC modulation in the NS-ROI and the VSD response in the pre-and post-ROIs.

Finally, we wanted to compare the MS modulation in the blank trials with the MS modulation in the NS-ROI of the stimulated trials (Fig. 4D). The amplitude and dynamics of the MS modulation in both conditions is highly similar as indicated by the value of the Pearson correlation (r=0.96, p<0.001). In summary, the MS modulation in blank trials, appears also in the non-stimulated V1 region, during visual stimulation.

### The MS modulation cannot be explained by an external motion of the fixation point

Thus far, we have studied V1 population responses to MSs generated by the animals, spontaneously, while the monkeys were maintaining fixation around a fixation point (FP). To test whether the MSs modulation might be explained by the FP movement following a MS, we performed a set of control experiments on a third monkey (monkey B). Here we artificially moved the FP or a tiny point located at the perimeter of the FP (i.e at the surrounding border of the FP; see Materials and Methods). Figure 5A shows the experimental design of the control experiment: the FP (or a small dot located at the perimeter of the FP), moved inward, into the imaging chamber or outward, i.e. away from the imaging chamber. The inward motion of the FP simulated a MS away from the imaging chamber and the outward motion of the FP stimulated a MS towards the imaging chamber (see Materials and Methods). To mimic real MSs properties, we used similar MSs amplitude and MSs frequency: the points shifted with an amplitude of 0.3-0.6 deg, inward or outward of the imaging chamber, for 200-300 ms and then returned back to the initial position.

**Figure 5:**
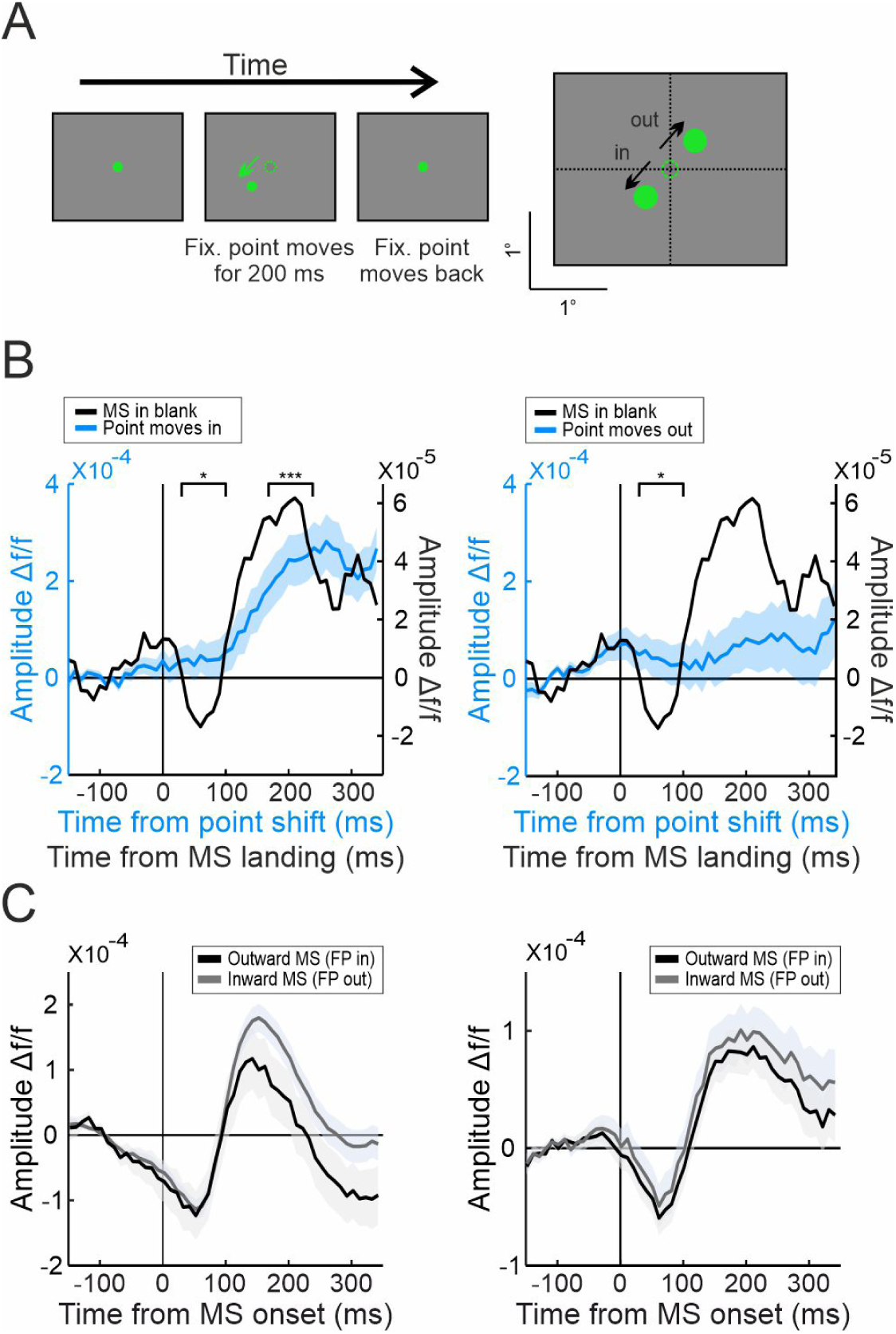
Artificial movement of the fixation point in comparison with the microsaccade modulation. **(A)** A schematic illustration of the control paradigm with an artificial movement of the fixation point (FP). Left: while the animal is fixating, the FP is shifted over the screen (in this example by ∼0.3 deg), mimicking a typical MS amplitude. Following 200-300 ms the FP returned to its initial point. Right: Movement direction of the FP: inward or outward of the imaging chamber. **(B)** Left: a comparison between the V1 VSD response to a MS (black curve, right y-axis; t=0 is the MS landing; n=59 MSs) and the response generated by an artificial inward movement of the FP (cyan curve, left y axis; t=0 is movement onset; n=30 trials). Top bars with asterisk depict the time windows with significant difference between the two responses. Right: same as for left, but when the FP moves outwards (n=26 trials). Baseline activity was subtracted from each condition (Wilcoxon rank sum; * p > 0.05; *** p > 0.001). **(C)** The MS modulation divided into two classes: MSs directed inward (leading to FP out of the chamber; gray curve) or MS outward (leading to FP into the chamber; black curve) of the chamber for each monkey. Left: monkey L, MSs with inward direction (n=52), MSs with outward direction (n=167). Right: monkey G, MSs with inward direction (n=146), MSs with outward direction (n=107). The correlation coefficient (r) between the MS modulation for the two MS directions: r=1, p<0.001, for both monkeys. Shaded area represents ±1SEM across MSs.

Figure 5B shows the VSD TC of response in V1 for the two control conditions (t=0 for FP motion, cyan curves, left y-axes): inward motion (Fig. 5B left) and outward motion (Fig. 5B right).

For comparison purposes, in the same graphs, we plotted the MS modulation in the blank trials of the same animal, aligned on MS landing (t=0, black curve; right y-axes; the two y-axes have different scales because the V1 response to the FP movement was much larger than the MS modulation in the blank trials). Figure 5B left shows that following an inward movement of the FP (the FP moved into the imaging chamber), a very sluggish and slow response gradually develops in V1. This response may reflect low sub-threshold spread of activation due to the FP stimulation in the new location (note that this response is much lower and slower than a typical V1 response to a visual stimulus). Figure 5B right shows that following an outward motion of the FP (the FP moved away from the imaging chamber), there is very little change in V1 response. To quantify the response differences generated by the artificial FP movement versus the MS modulation (in the absence of visual stimulation) we compared the response amplitude during the MS suppression (30-100 ms after MS Landing) and at peak enhancement (170-240 ms after MS Landing) in the inward movement (Fig. 5B left). The response amplitude in the FP control stimulus during the time window of MS suppression (30-100 ms) was 3.52±1.22×10^-5^ ΔF/F (n=30 trials), which was significantly higher than the negative peak response in the MS modulation: -0.77±0.69×10^-5^ ΔF/F (n=59 MSs; Wilcoxon ranksum test p<0.05). The response amplitude for the FP control stimulus at the peak time of MS enhancement (170-240 ms) was 21.66±1.93×10^-5^ ΔF/F (n=30 trials), which was significantly higher than the response amplitude in the MS modulation 5.74±0.83×10^-5^ ΔF/F (n=59 MSs; Wilcoxon ranksum test p<0.001). Finally, the response amplitude for the FP outward movement, at the MS suppression phase (30-100 ms) was 4.5±1.42×10^-5^ ΔF/F (n=26 trials) that was significantly higher than the response amplitude in the MS modulation -0.77±0.69×10^-5^ ΔF/F (n=59 trials; Wilcoxon ranksum test p<0.05). These results suggest that the influences of artificial movement of the FP and the MS modulation are different.

The VSD responses in the FP control experiments showed small changes between the inward and outward FP movement. We therefore decided to test whether the direction of real MSs can induce different V1 response. The VSD response for inward MSs (shifting the FP outward the chamber, gray curve) and for outward MSs (shifting the FP inward, towards the chamber, black curve) is plotted in Fig. 5C for both monkeys. The VSD responses to both directions of real MSs had similar amplitude and dynamics and showed a high correlation between the TC for the two MSs directions (r=1, p < 0.001 for monkeys L and G). In summary, the MS modulation induced by real MSs is different from that generated by an artificial movement of the FP. Moreover, MSs with opposite directions, that shift the FP towards or outwards of the imaging chamber, generate a highly similar VSD response, in accordance with a recent study in V1 of monkeys (Wu et al., 2022).

## Discussion

Our vision appears as stable despite the incessant eye movements that are leading to continual movement of the visual input over the retina. This can suggest the existence of an extra-retinal neuronal signal that can be used for correcting the stimulus shift and lead to the generation of visual stability. The few neurophysiological studies that investigated this topic in MSs, did not focus on V1 fovea, the area that is expected to be most influenced by the tiny fixational saccades. Moreover, most of these studies focused mainly on MSs effects in the presence of a visual stimulus. In our work we implemented a different approach: using VSDI, we characterized the influence of MSs on V1 activity at the fovea, in the absence of visual stimulation, except for a small fixation point. Following MSs, we found a biphasic modulation in V1 population response. Additionally, there is an increase in synchronization following a MS, supporting the existence of a common input into V1 that drives the neuronal population upon the MS onset. The biphasic modulation could not be explained by an artificial movement of the FP and co-existed in V1 next to the MS modulation in the presence of a visual stimulus.

### MSs effect in V1 reveal a biphasic modulation

Some of the studies that investigated the influence of fast eye-movement in the presence of a visual stimulus, suggested that the modulation in V1 can be explained by the stimulus shift over the retina: for saccades (Idrees et al., 2020; Parker et al., 2023) and for MSs (Gur and Snodderly, 1997; Leopold and Logothetis, 1998; Meirovithz et al., 2012; Gilad et al., 2017). However, other studies raised argumentations beyond the stimulus related component, and reported on the existence of an ERI to V1 (Kogan et al., 2008; Bremmer et al., 2009; Troncoso et al., 2015; Niemeyer et al., 2022; Wu et al., 2022). Yet, most of these studies were performed when a visual stimulus was displayed in the visual field, which makes the interpretation more complicated.

Following MS onset, we found a biphasic modulation in V1 population response, comprised of a suppression followed by enhancement. This biphasic modulation could not be explained by an artificial movement of the FP, thus further supporting the existence of an ERI to V1. Previous studies on saccades in the presence or absence of visual stimuli, reported on neural modulation that can support ERI in various visual cortical areas. This neural modulation showed typically suppression or/and enhancement following the saccade onset (Royal et al., 2006; Rajkai et al., 2008; Bremmer et al., 2009; Miura and Scanziani, 2022; Niemeyer et al., 2022; Denagamage et al., 2023). An evidence for ERI in MSs was reported by just a few studies (Snodderly et al., 2001; Troncoso et al., 2015; Wu et al., 2022), and some of them reported on a biphasic modulation comprised of suppression (70-100 ms from MS onset) followed by enhancement (160-200 ms from MS onset). These findings are in accordance with our results.

The suppression phase of the neural modulation in large saccades and fixational saccades was suggested to play a role in ’saccadic suppression’, a behavioral phenomenon characterized by a transient reduction in visual sensitivity around the saccadic onset (Volkmann, 1986; Diamond et al., 2000; Burr, 2004; Binda and Morrone, 2018) and recently also around MSs onset (Scholes et al., 2018; Intoy et al., 2021; but see also Martinez-Conde et al., 2013). Post-saccadic neural enhancement was suggested as a mechanism that can underly behavioral enhancement of visual processing after saccades, such as increase in visual sensitivity (Burr and Ross, 1982), increased contrast sensitivity (Diamond et al., 2000) and reduction in reaction times (Johns et al., 2009). Other neurophysiological studies reported that the post-saccadic neural enhancement is correlated with increased visual sensitivity to high spatial frequencies (Niemeyer et al., 2022), sharpening of orientation tuning curves (Ibbotson and Krekelberg, 2011) and faster evolvement of figure-ground segregation (Gilad et al., 2014). In recent years, behavioral enhancement of visual processing was also reported in relation to MSs, for example improved performance in visual acuity demanding tasks (Poletti et al., 2013; Ko et al., 2016; Intoy and Rucci, 2020) and enhanced sensitivity for a specific range of spatial frequencies 100-200 ms after MS onset (Bellet et al., 2017; Scholes et al., 2018; Intoy et al., 2021; but see: Mostofi et al., 2016). Finally, saccades and MSs were suggested to serve as a clock which parses vision between fixation epochs (Paradiso et al., 2012). According to this notion, there is a link between the two behavioral phenomena of saccadic suppression and the followed increased visual enhancement: their temporal sequence allows to actively mask the pre-saccadic stimulus and boost the response to the post-saccadic stimulus (Wurtz, 2008; Ibbotson and Krekelberg, 2011; Paradiso et al., 2012; Binda and Morrone, 2018).

### Mirosaccades induce increased synchronization

The neuronal population in V1 showed an increase in synchronization shortly after MS onset, the peak amplitude of the synchronization occurred ∼ 100 ms after MS onset. Previous studies of neuronal synchronization in relation to saccades and MSs in the visual cortex, were reported in animals that were presented with various visual stimuli (Leopold and Logothetis, 1998; Martinez-Conde et al., 2004; Maldonado et al., 2008; Bosman et al., 2009; Meirovithz et al., 2012; Lowet et al., 2016; Niemeyer et al., 2022). A previous VSD study in fixating monkeys, reported that MSs performed during the presence of small and large visual stimuli, induced increased synchronization in V1 approximately 100 ms after MS onset (Meirovithz et al., 2012). Bosman et al., 2009 reported on increased coherence in V1 and V4 following MSs and Lowet et al., 2018 reported on gamma band synchronization in V1 and V2. These observations were suggested to be linked with enhanced information processing of the stimulus, in its new landing position, following the spatial shift over the retina (Meirovithz et al., 2012; Niemeyer et al., 2022).

The increase in synchronization following a MS onset, that we found, in the absence of visual stimulation can be linked to the above reports and may reflect the neural preparation in V1 for enhanced processing of an upcoming stimulus. In addition, Martinez-Conde et al., 2000, 2002 suggested that the synchronization after each MS allows the MS signal to propagate and synchronize the neural activity across the visual areas.

### Possible neural sources for the MS modulation

The biphasic MS modulation in our work, could not be explained by an artificial movement of the FP, which further supports the existence of an ERI to V1. The few neurophysiological studies which reported on evidence for ERI to V1 in MSs, suggested several possible neuronal sources: proprioceptive signals, corollary discharge (efference copy of the motor command to the oculomotor muscles), global motion signals and attentional signals (Kogan et al., 2008; Troncoso et al., 2015; Wu et al., 2022). Studies which focused on saccades offered similar sources of ERI signals to the visual system (Hopkins Duffy and Lee Burchfiel, 1975; Corbetta et al., 1998; Royal et al., 2006; Rajkai et al., 2008; Ibbotson and Krekelberg, 2011; Miura and Scanziani, 2022; Niemeyer et al., 2022).

The superior colliculus (SC) plays a key role in motor execution of saccades and MSs (Hafed et al., 2009). The corollary discharge (CD) refers to a copy of the efference motor command to the oculomotor muscles. A CD signal, starting tens of ms before MSs onset, can emerge from the intermediate layers of the SC and arrive to the visual cortex via different pathways (Wurtz, 2008; Krauzlis et al., 2017). A recent study by Denagamage et al., 2023 reported on neural suppression following saccades, that initiated by the activation of inhibitory neurons in layer IV in area V4. The authors suggested the Pulvinar as a possible neural source for this response, which was previously suggested to carry CD signal from the SC to various visual areas such as MT in primates and V1 in rodents and primates (Stepniewska et al., 2000; Shipp, 2004; Berman and Wurtz, 2010; Kuang et al., 2012; Schneider et al., 2020, 2023; Miura and Scanziani, 2022). Therefore, the initial suppression phase we report can fit to a similar possible pathway from the SC through the Pulvinar into the inhibitory neurons in V1. Another possible pathway of the CD travels through the mediodorsal nucleus of the thalamus to the frontal-eye-field (FEF; Sommer and Wurtz, 2002) and from there to V1 (through V4). However, this track might be too slow to account for the early neural suppression immediately after MS onset that can start even before the MS (Fig. 2C).

Proprioceptive signals from the oculomotor muscles can carry information about the eye position and movement kinematics. While such signals were previously shown to affect the neural activity in V1 (Trotter et al., 1993; Buisseret, 1995), most of the previous studies indicated that the oculomotor proprioceptive signal is too slow to account for the fast dynamics of the MS suppression (Balslev et al., 2012; Sun and Goldberg, 2016). One may speculate that a proprioceptive signal can be a relevant candidate for the late enhancement phase present in our results, yet the anatomical pathways that carry such information to the visual cortex were not reported yet.

Attention has been extensively linked with saccades and MSs (Corbetta et al., 1998; Hafed and Clark, 2002; Engbert and Kliegl, 2003; Martinez-Conde et al., 2004; Shipp, 2004). A well-established pathway initiates from FEF to V4 which mediates an attention-related input to V1 (Corbetta et al., 1998; Fries et al., 2001; Martinez-Conde et al., 2013; Denagamage et al., 2023).

Alongside, the SC and the Pulvinar might be also involved in a parallel pathway mediating the attentional signal from the FEF through these subcortical centers (Shipp, 2004). The attentional effects in V1 activity appear around 200-250 ms after stimulus onset (Roelfsema et al., 1998; Lamme and Roelfsema, 2000). This timing fits well with the temporal dynamics of the enhancement in the MS modulation, as we reported.

### The relation between MS modulation in the presence and absence of a visual stimulus

Our study showed the co-existence of two different MS modulations (Fig. 4): one that is induced by the visual stimulus shift over the retina (visually driven modulation), and another that is evident in the non-stimulated region. The visually related modulation is ten times larger than the MS modulation in the non-stimulated region and the latter is highly similar to the MS modulation in the blank trials. Interestingly, the temporal evolvement of the MS modulation in the two conditions are different. The suppression phase in non-stimulated region (and MS modulation in the blank) occurs before the activation decrease for the visually driven modulation (pre-MS ROI). In contrast, the enhancement phase in the non-stimulated region (and MS modulation in the blank) is delayed relative to the response increase in the post-MS ROI. This may further support the notion that sources for the two modulations are, at least partially, independent, and each carries different information. Additional future studies are required to uncover and dissect the anatomical pathways underlying the MS modulation in the absence of visual stimulation and how these interact with visual stimulation and possibly leads to the production of visual stabilization.

## References

1. Ahissar E, Arieli A, Fried M, Bonneh Y (2016) On the possible roles of microsaccades and drifts in visual perception. Vision Res 118:25–30.

2. Arieli A, Grinvald A, Slovin H (2002) Dural substitute for long-term imaging of cortical activity in behaving monkeys and its clinical implications. J Neurosci Methods 114:119–133.

3. Ayzenshtat I, Meirovithz E, Edelman H, Werner-Reiss U, Bienenstock E, Abeles M, Slovin H (2010) Precise spatiotemporal patterns among visual cortical areas and their relation to visual stimulus processing. J Neurosci 30:11232–11245.

4. Bach M, Bouis D, Fischer B (1983) An accurate and linear infrared oculometer. J Neurosci Methods 9:9–14.

5. Balslev D, Himmelbach M, Karnath H-O, Borchers S, Odoj B (2012) Eye Proprioception Used for Visual Localization Only If in Conflict with the Oculomotor Plan. J Neurosci 32:8569 LP – 8573.

6. Bellet J, Chen CY, Hafed ZM (2017) Sequential hemifield gating of α-and β-behavioral performance oscillations after microsaccades. J Neurophysiol 118:2789–2805.

7. Berman RA, Wurtz RH (2010) Functional identification of a pulvinar path from superior colliculus to cortical area MT. J Neurosci 30:6342–6354.

8. Binda P, Morrone MC (2018) Vision during saccadic eye movements. Annu Rev Vis Sci 4:193– 213.

9. Bosman CA, Womelsdorf T, Desimone R, Fries P (2009) A microsaccadic rhythm modulates gamma-band synchronization and behavior. J Neurosci 29:9471–9480.

10. Bremmer F, Kubischik M, Hoffmann KP, Krekelberg B (2009) Neural dynamics of saccadic suppression. J Neurosci 29:12374–12383.

11. Buisseret P (1995) Influence of extraocular muscle proprioception on vision. Physiol Rev 75:323– 338.

12. Burr DC, Ross J (1982) Contrast sensitivity at high velocities. Vision Res 22:479–484.

13. Burr M (2004) Visual perception during saccades. In: The visual neurosciences., vol 2. (Cook, J. E., Chalupa, L. M., & Werner JS, ed), pp 1391–1401. Cambridge, Massachusetts: MIT Press.

14. Chen CY, Hafed ZM (2013) Postmicrosaccadic enhancement of slow eye movements. Ann Intern Med 158:5375–5386.

15. Chen CY, Hoffmann KP, Distler C, Hafed ZM (2019) The Foveal Visual Representation of the Primate Superior Colliculus. Curr Biol 29:2109–2119.e7.

16. Cherici C, Kuang X, Poletti M, Rucci M (2012) Precision of sustained fixation in trained and untrained observers. J Vis 12:1–16.

17. Corbetta M, Akbudak E, Conturo TE, Snyder AZ, Ollinger JM, Drury HA, Linenweber MR, Petersen SE, Raichle ME, Van Essen DC, Shulman GL (1998) A common network of functional areas for attention and eye movements. Neuron 21:761–773.

18. Denagamage S, Morton MP, Hudson N V., Reynolds JH, Jadi MP, Nandy AS (2023) Laminar mechanisms of saccadic suppression in primate visual cortex. Cell Rep 42:112720.

19. Diamond MR, Ross J, Morrone MC (2000) Extraretinal Control of Saccadic Suppression. J Neurosci 20:3449–3455.

20. Engbert R, Kliegl R (2003) Microsaccades uncover the orientation of covert attention. Vision Res 43:1035–1045.

21. Engbert R, Mergenthaler K (2006) Microsaccades are triggered by low retinal image slip. Proc Natl Acad Sci U S A 103:7192–7197.

22. Fries P, Reynolds JH, Rorie AE, Desimone R (2001) Modulation of oscillatory neuronal synchronization by selective visual attention. Science 291:1560–1563.

23. Gilad A, Oz R, Slovin H (2017) Uncovering the Spatial Profile of Contour Integration from Fixational Saccades: Evidence for Widespread Processing in V1. Cereb Cortex 27:5261–5273.

24. Gilad A, Pesoa Y, Ayzenshtat I, Slovin H (2014) Figure-ground processing during fixational saccades in V1: Indication for higher-order stability. J Neurosci 34:3247–3252.

25. Green DG (1970) Regional variations in the visual acuity for interference fringes on the retina. J Physiol 207:351–356.

26. Grinvald A, Hildesheim R (2004) VSDI: a new era in functional imaging of cortical dynamics. Nat Rev Neurosci 5:874–885.

27. Gur M, Snodderly DM (1997) Visual Receptive Fields of Neurons in Primary Visual Cortex (VI) Move in Space with the Eye Movements of Fixation. 37:257–265.

28. Hafed ZM, Clark JJ (2002) Microsaccades as an overt measure of covert attention shifts. Vision Res 42:2533–2545.

29. Hafed ZM, Goffart L, Krauzlis RJ (2009) A Neural Mechanism for Microsaccade Generation in the Primate Superior Colliculus. Science 323:940–943.

30. Hopkins Duffy F, Lee Burchfiel J (1975) Eye movement-related inhibition of primate visual neurons. Brain Res 89:121–132.

31. Ibbotson M, Krekelberg B (2011) Visual perception and saccadic eye movements. Curr Opin Neurobiol 21:553–558.

32. Idrees S, Baumann MP, Franke F, Münch TA, Hafed ZM (2020) Perceptual saccadic suppression starts in the retina. Nat Commun 11:1–19.

33. Intoy J, Mostofi N, Rucci M (2021) Fast and nonuniform dynamics of perisaccadic vision in the central fovea. Proc Natl Acad Sci U S A 118.

34. Intoy J, Rucci M (2020) Finely tuned eye movements enhance visual acuity. Nat Commun 11:1– 11.

35. Jancke D, Chavane F, Naaman S, Grinvald A (2004) Imaging cortical correlates of illusion in early visual cortex. Nature 428:423–426.

36. Johns M, Crowley K, Chapman R, Tucker A, Hocking C (2009) The effect of blinks and saccadic eye movements on visual reaction times. Attention, Perception, Psychophys 71:783–788.

37. Ko H kyoung, Snodderly DM, Poletti M (2016) Eye movements between saccades: Measuring ocular drift and tremor. Vision Res 122:93–104.

38. Kogan I, Gur M, Snodderly DM (2008) Saccades and drifts differentially modulate neuronal activity in V1: Effects of retinal image motion, position, and extraretinal influences. J Vis 8:1–25.

39. Krauzlis RJ, Goffart L, Hafed ZM (2017) Neuronal control of fixation and fixational eye movements. Philos Trans R Soc B Biol Sci 372.

40. Kuang X, Poletti M, Victor JD, Rucci M (2012) Temporal encoding of spatial information during active visual fixation. Curr Biol 22:510–514.

41. Lamme VA, Roelfsema PR (2000) The distinct modes of vision offered by feedforward and recurrent processing. Trends Neurosci 23:571–579.

42. Leopold DA, Logothetis NK (1998) Microsaccades differentially modulate neural activity in the striate and extrastriate visual cortex. Exp Brain Res 123:341–345.

43. Lowet E, Gips B, Roberts MJ, De Weerd P, Jensen O, van der Eerden J (2018) Microsaccade-rhythmic modulation of neural synchronization and coding within and across cortical areas V1 and V2.

44. Lowet E, Roberts MJ, Bosman CA, Fries P, de Weerd P (2016) Areas V1 and V2 show microsaccade-related 3-4-Hz covariation in gamma power and frequency. Eur J Neurosci 43:1286–1296.

45. Maldonado P, Babul C, Singer W, Rodriguez E, Berger D, Grün S (2008) Synchronization of neuronal responses in primary visual cortex of monkeys viewing natural images. J Neurophysiol 100:1523–1532.

46. Martinez-Conde S, Macknik SL, Hubel DH (2000) Microsaccadic eye movements and firing of single cells in the striate cortex of macaque monkeys. Nat Neurosci 3:251–258.

47. Martinez-Conde S, Macknik SL, Hubel DH (2002) The function of bursts of spikes during visual fixation in the awake primate lateral geniculate nucleus and primary visual cortex. Proc Natl Acad Sci U S A 99:13920–13925.

48. Martinez-Conde S, Macknik SL, Hubel DH (2004) The role of fixational eye movements in visual perception. Nat Rev Neurosci 5:229–240.

49. Martinez-Conde S, Macknik SL, Troncoso XG, Hubel DH (2009) Microsaccades: a neurophysiological analysis. Trends Neurosci 32:463–475.

50. Martinez-Conde S, Otero-Millan J, MacKnik SL (2013) The impact of microsaccades on vision: Towards a unified theory of saccadic function. Nat Rev Neurosci 14:83–96.

51. Meirovithz E, Ayzenshtat I, Bonneh YS, Itzhack R, Werner-Reiss U, Slovin H (2010) Population response to contextual influences in the primary visual cortex. Cereb Cortex 20:1293–1304.

52. Meirovithz E, Ayzenshtat I, Werner-Reiss U, Shamir I, Slovin H (2012) Spatiotemporal effects of microsaccades on population activity in the visual cortex of monkeys during fixation. Cereb Cortex 22:294–307.

53. Miura SK, Scanziani M (2022) Distinguishing externally from saccade-induced motion in visual cortex. Nature 610:135–142.

54. Moiseenko GA, Vakhrameeva OA, Lamminpiya AM, Pronin S V., Maltsev DS, Sukhinin M V., Vershinina EA, Kovalskaya AA, Koskin SA, Shelepin YE (2018) Dependence between the Size of the Foveola and the Parameters of Visual Perception. Hum Physiol 44:510–516.

55. Mostofi N, Boi M, Rucci M (2016) Are the visual transients from microsaccades helpful? Measuring the influences of small saccades on contrast sensitivity. Vision Res 118:60–69.

56. Niemeyer JE, Akers-Campbell S, Gregoire A, Paradiso MA (2022) Perceptual enhancement and suppression correlate with V1 neural activity during active sensing. Curr Biol 32:2654–2667.e4.

57. Paradiso MA, Meshi D, Pisarcik J, Levine S (2012) Eye movements reset visual perception. J Vis 12:11.

58. Parker PRL, Martins DM, Leonard ESP, Casey NM, Sharp SL, Abe ETT, Smear MC, Yates JL, Mitchell JF, Niell CM (2023) A dynamic sequence of visual processing initiated by gaze shifts. Nat Neurosci 26:2192–2202.

59. Poletti M, Listorti C, Rucci M (2013) Microscopic eye movements compensate for nonhomogeneous vision within the fovea. Curr Biol 23:1691–1695.

60. Rajkai C, Lakatos P, Chen C-M, Pincze Z, Karmos G, Schroeder CE (2008) Transient cortical excitation at the onset of visual fixation. Cereb Cortex 18:200–209.

61. Roelfsema PR, Lamme VA, Spekreijse H (1998) Object-based attention in the primary visual cortex of the macaque monkey. Nature 395:376–381.

62. Rolfs M (2009) Microsaccades: Small steps on a long way. Vision Res 49:2415–2441.

63. Ross J, Morrone MC, Goldberg ME, Burr DC (2001) Changes in visual perception at the time of saccades. Trends Neurosci 24:113–121.

64. Royal DW, Sáry G, Schall JD, Casagrande VA (2006) Correlates of motor planning and postsaccadic fixation in the macaque monkey lateral geniculate nucleus. Exp Brain Res 168:62– 75.

65. Rucci M, Victor JD (2015) The unsteady eye: An information-processing stage, not a bug. Trends Neurosci 38:195–206.

66. Schneider L, Dominguez-Vargas AU, Gibson L, Kagan I, Wilke M (2020) Eye position signals in the dorsal pulvinar during fixation and goal-directed saccades. J Neurophysiol 123:367–391.

67. Schneider L, Dominguez-Vargas AU, Gibson L, Wilke M, Kagan I (2023) Visual, delay, and oculomotor timing and tuning in macaque dorsal pulvinar during instructed and free choice memory saccades. Cereb Cortex 33:10877–10900.

68. Scholes C, McGraw P V., Roach NW (2018) Selective modulation of visual sensitivity during fixation. J Neurophysiol 119:2059–2067.

69. Shipp S (2004) The brain circuitry of attention. Trends Cogn Sci 8:223–230.

70. Shoham D, Glaser DE, Arieli A, Kenet T, Wijnbergen C, Toledo Y, Hildesheim R, Grinvald A (1999) Imaging cortical dynamics at high spatial and temporal resolution with novel blue voltage-sensitive dyes. Neuron 24:791–802.

71. Shtoyerman E, Arieli A, Slovin H, Vanzetta I, Grinvald A (2000) Long-term optical imaging and spectroscopy reveal mechanisms underlying the intrinsic signal and stability of cortical maps in V1 of behaving monkeys. J Neurosci 20:8111–8121.

72. Slovin H, Arieli A, Hildesheim R, Grinvald A (2002) Long-term voltage-sensitive dye imaging reveals cortical dynamics in behaving monkeys. J Neurophysiol 88:3421–3438.

73. Snodderly DM, Kagan I, Moshe G (2001) Selective activation of visual cortex neurons by fixational eye movements: Implications for neural coding. Vis Neurosci 18:259–277.

74. Sommer MA, Wurtz RH (2002) A pathway in primate brain for internal monitoring of movements. Science 296:1480–1482.

75. Stepniewska I, Qi HX, Kaas JH (2000) Projections of the superior colliculus to subdivisions of the inferior pulvinar in New World and Old World monkeys. Vis Neurosci 17:529–549.

76. Sun LD, Goldberg ME (2016) Corollary Discharge and Oculomotor Proprioception: Cortical Mechanisms for Spatially Accurate Vision. Annu Rev Vis Sci 2:61–84.

77. Troncoso XG, McCamy MB, Jazi AN, Cui J, Otero-Millan J, MacKnik SL, Costela FM, Martinez-Conde S (2015) V1 neurons respond differently to object motion versus motion from eye movements. Nat Commun 6:1–10.

78. Trotter Y, Celebrini S, Beaux JC, Grandjean B, Imbert M (1993) Long-term dysfunctions of neural stereoscopic mechanisms after unilateral extraocular muscle proprioceptive deafferentation. J Neurophysiol 69:1513–1529.

79. Volkmann FC (1986) Human visual suppression. Vision Res 26:1401–1416.

80. Wu Y, Wang T, Zhou T, Li Y, Yang Y, Dai W, Zhang Y, Han C, Xing D (2022) V1-bypassing suppression leads to direction-specific microsaccade modulation in visual coding and perception. Nat Commun 13:1–14.

81. Wurtz RH (2008) Neuronal mechanisms of visual stability. Vision Res 48:2070–2089.

